# Low birth weight in Bangladesh and associated maternal and socioeconomic factors: results from a recent nationally-representative survey

**DOI:** 10.1101/546333

**Authors:** Fariha Binte Hossain, Gourab Adhikary, Yasir Arafat, Md Shajedur Rahman Shawon

**Affiliations:** Independent Researcher, 368/3D, Ahmednagar, Mirpur-1, Dhaka 1216, Bangladesh.; Independent Researcher, Peace Tower, 8/3 Ponditpara, Mymensingh; Department of Health and Nutrition, Save the Children Bangladesh, House No. CWN (A) 35, Road No. 43 Gulshan-2, Dhaka 1212, Bangladesh.; Nuffield Department of Population Health, University of Oxford, OX3 7LF, Oxford, UK.

**Keywords:** Low birth weight, socioeconomic factor, ANC, Bangladesh, DHS

## Abstract

**Objectives:** Little is known about the relative contributions of maternal and socioeconomic factors on low birth weight in Bangladesh and whether they differ by sex. We examined the prevalence and associated maternal and socioeconomic factors of low birth weight separately among boys and girls.

**Design and settings:** This is a cross-sectional study based on the Bangladesh Demographic and Health Survey 2014.

**Participants:** A total of 4728 children with information on birth size were included in this study.

**Outcome measure:** Low birth weight was defined according to mother’s perception of birth size of their children. Logistic regression analysis was used to estimate the association between maternal and socioeconomic factors with low birth weight.

**Results:** The overall prevalence of low birth weight was 17.8% among boys and 22.4% among girls. Low birth weight was associated with maternal factors like maternal age of <20 years at birth (adjusted OR vs. 20-29 years: 1.40, 95% CI 1.09-1.78), and maternal undernutrition (adjusted OR 1.33, 95% CI 1.05-1.69) among boys while only the association with maternal undernutrition was significant among girls. The association for no antenatal care visit was explained by socioeconomic factors. Lower level of mother’s education and poorest wealth index were found to be associated with low birth weight in both sexes.

**Conclusion:** Our study identifies that maternal factors are associated with increased risk of having low birth weight babies, which cannot be explained by socioeconomic factors, and vice versa. Community-based interventions to reduce low birth weight in Bangladesh should focus on these factors.

**Strengths and limitations of this study:**

- To the best of our knowledge our study is the first study in Bangladesh that has looked at the factors for low birth weight separately among boys and girls.
- We used multiple logistic regression to examine the associations of maternal and socioeconomic factors with low birth weight in a large and nationally-representative sample.
- This study is limited because we used mother’s perception of child’s size at birth to define low birth weight.
- We did not have information about gestational age and thereby could not take prematurity into account in our study.

## INTRODUCTION

Low birth weight is a major global health problem because it affects a large number of babies worldwide and is associated with a wide range of short- and long-term consequences on morbidity and mortality [1,2]. According to the World Health Organization (WHO), birth weight of less than 2500 grams, irrespective of gestational age, is considered as low birth weight [1]. Recent estimates suggest that approximately 15% of 20% of all babies born in a year suffer from low birth weight [3]. There is a large disparity in the burden of low birth weight in the developed and developing countries. Also, south Asian countries harbour a large number of low birth weight babies due to higher birth rate as well as higher prevalence rate of low birth weight [1,4].

Low birth weight is associated with higher rates of diarrhoea, respiratory illness, hospitalisation, and eventually mortality during infancy and early childhood [5]. Recent evidences also suggest that low birth weight has long-term consequences on adult chronic diseases like diabetes, hypertension, cardiovascular diseases [6]. Therefore, tackling low birth weight has been considered as a priority area in global health and development.

Research efforts have been made previously to identify the burden and determinants of low birth weight in Bangladesh, but little is known about the possible sex differences in them. Also, it is not clear whether the relative contributions of maternal factors can be explained by socioeconomic inequalities, and vice versa. In this study, we examined the prevalence and associated maternal and socioeconomic factors of low birth weight among boys and girls, using a recent nationally-representative survey in Bangladesh.

## METHODS

### Study design and data sources

This is a cross-sectional study based on the Bangladesh Demographic and Health Survey (BDHS) 2014. Details about BDHS has been described elsewhere [7] and summarized below. At every 3-5 years, BDHS surveys are conducted by the National Institute for Population Research and Training (NIPORT) of the Ministry of Health and Family Welfare, Government of Bangladesh with technical assistance from the ICF International, located in Calverton, Maryland, USA. BDHS surveys are nationally representative surveys and are based on two-stage stratified sampling of households. In the first stage, census enumeration areas are selected using probability proportional to size (PPS) sampling technique through statistics provided by the Bangladesh Bureau of Statistics (BBS). In the second stage, households are selected through systematic random sampling from the complete listing of households within a selected enumeration area [7].

Ethical approval for BDHS 2014 was taken from the ICF International Institutional Review Board. All participating gave Informed consent. The data files are freely available from the MEASURE Demographic and Health Surveys (DHS) website (www.dhsprogram.com). We received authorization from the DHS program for using the relevant dataset for this analysis. The data we received were anonymized for protection of privacy, anonymity and confidentiality.

This survey had a response rate of 98%. The detailed questionnaires of this survey are available in the final report of BDHS 2014. We included those children who were born in the last three years before the survey and had information on their size at birth.

### Low birth weight

In BDHS 2014, mothers were asked about the perceived size at birth of their children born in the three years preceding the survey. The available responses were very large, larger than average, average, smaller than average, and very small at birth. For our analysis, we categorized those with responses very small and smaller than average as low birth weight babies, and those with very large, larger than average, and average as normal weight babies. BDHS 2014 did not have information on birth weight and therefore, we could not define low birth weight based on conventional cut-off of less than 2500 grams. However, a previous validation study using DHS data from Cambodia, Kazakhstan and Malawi suggested that mother’s perception of baby’s size at birth agreed well with recorded birth weight and could be a useful tool to obtain more accurate estimates of low birth weight [8].

### Child, maternal, and socioeconomic variables

BDHS 2014 collected information about the characteristics of selected households and their respondents by using face-to-face interview with standardized questionnaires, taken by trained personnel. In this study, we grouped the available explanatory variables into three groups: child, maternal, and socioeconomic. Child variables were age, sex, and birth order of the child. Maternal factors were mother’s age at birth, maternal undernutrition, parity, number of antenatal care (ANC) visits. Socioeconomic variables included area of residence, mother’s education level, and household wealth index.

Mother’s age at birth was categorized into three groups: <20 years, 20-29 years and ≥30 years. We also categorized number of ANC visits into three groups: no visit, 1–3 visits, and ≥4 visits. Parity was dichotomized (i.e. 0-2, and ≥3) while maternal undernutrition was defined based on body mass index (BMI) <18.5 kg/m^2^. BMI was calculated based on the measured height and weight at the time of survey. Place of residence (rural and urban) was defined according to country-specific definitions. Mother’s education was grouped into no education, primary, secondary, and higher. For household’s wealth index, BBS constructed a country-specific index using principal components analysis from data on household assets including durable goods (i.e. bicycles, televisions etc.) and dwelling characteristics (i.e. sanitation, source of drinking water and construction material of house etc.) [7]. This wealth index was then categorized into five groups (i.e. poorest, poorer, middle, richer, and richest) based on the quintile distribution of the sample.

### Statistical analysis

All of our analyses were stratified by sex of the child. We first looked at the percent distribution of all included child, maternal, and socioeconomic characteristics of the participants. For calculating prevalence of low birth weight, we followed the DHS guide’s instructions for analysis – we used the sampling weights given in the dataset to yield estimates which were representative of the Bangladeshi population [9]. 95% confidence intervals (CIs) for prevalence estimates were calculated using a logit transform of the estimate. We then used simple logistic regressions to estimate crude odds ratios (ORs) for low birth weight with 95% CIs, by maternal and socioeconomic factors. We also estimated adjusted ORs for birth weight using multiple logistic regression by adjusting for child, maternal, and socioeconomic variables, as required.

All analyses were performed using Stata v14.2 (Statacorp, College Station, TX, USA). Considering the two-stage stratified cluster sampling in BDHS 2014, we applied Stata’s survey estimation procedures (*“svy”* command) for regression analyses.

## RESULTS

### Participants’ characteristics

Table 1 shows the percent distribution of all characteristics of the participants. A total of 4728 children (55% male) had valid information on size at birth. They had equal distribution for current age while most (60%) of them had birth order of two or more. At the time of child’s birth, almost two-thirds of the mothers were aged between 20-29 years. One in every four mothers were underweight. Only about one-third of all mothers had four or more ANC visits while 21% had no visit. Most (∼75%) of the included children were from rural area. Almost half of the mothers had secondary education and household wealth index was distributed in almost equal five groups. There were no significant differences between male and female children for these characteristics (Table 1).

**Table 1:** Percent distribution of all characteristics of the participants.

|  | Male n (%) | Female n (%) |
| --- | --- | --- |
| Number of children | 2575 | 2322 |
| Child's current age |  |  |
| 0 year | 809 (32.7) | 704 (31.5) |
| 1 year | 861 (34.8) | 771 (34.5) |
| 2 year | 802 (32.4) | 760 (33.9) |
| Child's birth order |  |  |
| 1 | 1033 (40.6) | 943 (40.6) |
| 2 or more | 1541 (59.9) | 1378 (59.4) |
| Mother's age at birth |  |  |
| <20 years | 704 (28.5) | 614 (27.5) |
| 20-29 years | 1393 (56.3) | 1269 (56.8) |
| ≥30 years | 375 (15.2) | 351 (15.7) |
| Parity |  |  |
| 0-2 | 1909 (74.2) | 1689 (72.7) |
| ≥3 | 665 (25.9) | 633 (27.3) |
| Maternal undernutrition (BMI <18.5 kg/m<sup>2</sup>) |  |  |
| Yes | 625 (24.4) | 577 (24.9) |
| No | 1944 (75.7) | 1744 (75.1) |
| ANC visit |  |  |
| 0 | 518 (21.3) | 473 (21.7) |
| 1-3 | 1135 (46.6) | 1052 (48.1) |
| 4 or more | 782 (32.1) | 660 (30.2) |
| Area of residence |  |  |
| Urban | 671 (26.0) | 593 (25.6) |
| Rural | 1903 (73.9) | 1728 (74.4) |
| Mother's education |  |  |
| No education | 371 (14.4) | 329 (14.2) |
| Primary | 706 (27.4) | 671 (28.9) |
| Secondary | 1235 (47.9) | 1095 (47.1) |
| Higher | 263 (10.2) | 227 (9.8) |
| Household's wealth index |  |  |
| Poorest | 554 (21.5) | 529 (22.8) |
| Poorer | 484 (18.8) | 447 (19.3) |
| Middle | 500 (19.4) | 439 (18.9) |
| Richer | 535 (20.8) | 460 (19.8) |
| Richest | 502 (19.5) | 447 (19.3) |

### Prevalence of low birth weight

The overall prevalence of low birth weight in Bangladesh was 17.8% and 22.4% for male and female children, respectively (Figure 1). Prevalence of low birth weight was particularly higher, irrespective of child’s sex, if mother’s age was less than 20 years at the time of child’s birth, mother had no ANC visit, and mother was undernourished (Table 2). There was no significant difference in low birth weight prevalence by parity.

**Table 2:** Prevalence of low birth weight by maternal factors, and their associations.

|  | Male |  |  | Female |  |  |
| --- | --- | --- | --- | --- | --- | --- |
|  | Prevalence <sup>a</sup> | Crude OR | Adjusted OR <sup>b</sup> | Prevalence <sup>a</sup> | Crude OR | Adjusted OR <sup>b</sup> |
| Mother's age at birth |  |  |  |  |  |  |
| <20 | 22.5 (19.7-24.7) | 1.38 (1.08-1.75) | 1.40 (1.09-1.78) | 22.8 (19.6-26.3) | 1.04 (0.82-1.33) | 1.07 (0.84-1.37) |
| 20-29 | 15.6 (13.7-17.6) | 1.00 (ref) | 1.00 (ref) | 20.7 (18.5-23.1) | 1.00 (ref) | 1.00 (ref) |
| ≥30 | 19.9 (16.6-23.8) | 1.24 (0.94-1.63) | 1.08 (0.79-1.48) | 22.3 (18.6-26.4) | 1.02 (0.78-1.35) | 0.79 (0.57-1.09) |
| ANC visit |  |  |  |  |  |  |
| 0 | 23.5 (20.1-27.4) | 1.62 (1.22-2.16) | 1.22 (0.89-1.70) | 27.9 (24.1-32.2) | 1.52 (1.14-2.01) | 1.23(0.89-1.68) |
| 1-3 | 16.7 (14.7-19.0) | 1.11 (0.86-1.43) | 0.98 (0.76-1.29) | 20.8 (18.4-23.3) | 1.10 (0.87-1.42) | 0.97 (0.74-1.26) |
| ≥4 | 14.9 (12.6-17.6) | 1.00 (ref) | 1.00 (ref) | 21.4 (18.4-24.7) | 1.00 (ref) | 1.00 (ref) |
| Maternal undernutrition (BMI<18.5 kg/m<sup>2</sup>) |  |  |  |  |  |  |
| No | 16.4 (14.8-18.1) | 1.00 (ref) | 1.00 (ref) | 20.4 (18.6-22.3) | 1.00 (ref) | 1.00 (ref) |
| Yes | 21.6 (8.6-25.0) | 1.43 (1.14-1.80) | 1.33 (1.05-1.69) | 28.2 (24.6-31.9) | 1.52 (1.22-1.89) | 1.40 (1.11-1.77) |
| Parity |  |  |  |  |  |  |
| 0-2 | 17.7 (16.0-19.5) | 1.00 (ref) | 1.00 (ref) | 23.0 (21.0-25.0) | 1.00 (ref) | 1.00 (ref) |
| ≥3 | 18.0 (15.3-21.1) | 1.07 (0.85-1.34) | 0.81 (0.63-1.05) | 20.7 (17.7-24.0) | 0.98 (0.78-1.23) | 0.84 (0.62-1.05) |

<sup>a</sup> Sampling weights were used to calculate prevalence
<sup>b</sup> Adjusted for child's current age, mother's education, area of residence and household's wealth index

**Figure 1:**
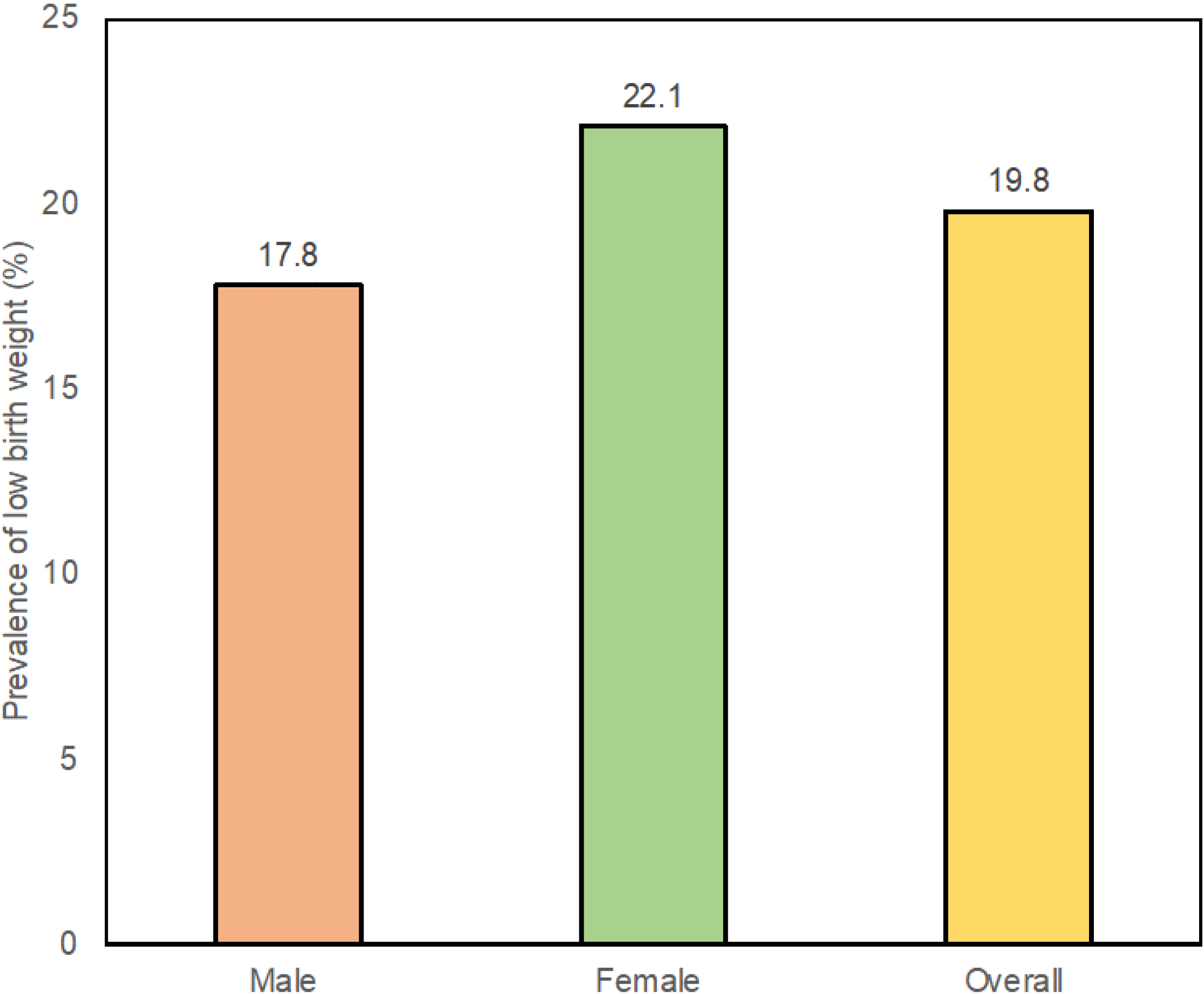
Prevalence of low birth weight in Bangladesh, overall and by sex.

Table 3 shows the prevalence of low birth weight by socioeconomic factors. There were significant differences in prevalence by maternal education and household wealth index. Mothers with no education had 26% of low birth weight both in male and female babies. Among mothers with higher education level there was a sex-difference in low birth weight prevalence (12% in male vs. 16% in female). Poorest households had 21% babies with low birth weight in males while 27% of female children had low birth weight in those households. There was no significant difference in low birth weight prevalence by area of residence.

**Table 3:** Prevalence of low birth weight by socioeconomic factors, and their associations.

|  | Male |  |  | Female |  |  |
| --- | --- | --- | --- | --- | --- | --- |
|  | Prevalence <sup>a</sup> | Crude OR | Adjusted OR <sup>b</sup> | Prevalence <sup>a</sup> | Crude OR | Adjusted OR <sup>b</sup> |
| Area of Residence |  |  |  |  |  |  |
| Urban | 15.9 (13.4-18.9) | 1.00 (Ref) | 1.00 (Ref) | 21.8 (18.6-25.3) | 1.00 (Ref) | 1.00 (Ref) |
| Rural | 18.4 (16.7-20.2) | 1.25 (0.99-1.56) | 1.15 (0.90-1.46) | 22.5 (20.6-24.5) | 1.18 (0.94-1.47) | 1.06 (0.83-1.35) |
| Mother's education |  |  |  |  |  |  |
| No education | 26.8 (22.5-31.6) | 2.89 (1.88-4.43) | 2.46 (1.49-4.05) | 26.2 (21.7-31.2) | 2.01 (1.31-3.09) | 1.97 (1.21-3.22) |
| Primary | 18.9 (16.3-22.0) | 1.87 (1.24-2.80) | 1.66 (1.06-2.60) | 23.2 (20.2-26.6) | 1.66 (1.12-2.44) | 1.52 (0.98-2.34) |
| Secondary | 15.6 (13.6-17.7) | 1.40 (0.95-2.08) | 1.36 (0.89-2.07) | 21.9 (19.5-24.5) | 1.35 (0.93-1.96) | 1.29 (0.86-1.94) |
| Higher | 12.4 (8.9-17.0) | 1.00 (Ref) | 1.00 (Ref) | 16.3 (11.9-21.7) | 1.00 (Ref) | 1.00 (Ref) |
| Household's wealth index |  |  |  |  |  |  |
| Poorest | 21.2 (17.9-24.8) | 1.92 (1.38-2.68) | 1.58 (1.07-2.32) | 26.5 (22.9-30.5) | 1.66 (1.21-2.27) | 1.42 (0.98-2.05) |
| Poorer | 17.8 (14.6-21.5) | 1.37 (0.96-1.96) | 1.24 (0.84-1.84) | 23.3 (19.6-27.4) | 1.33 (0.95-1.85) | 1.24 (0.86-1.80) |
| Middle | 18.7 (15.5-22.4) | 1.60 (1.13-2.27) | 1.53 (1.05-2.24) | 20.8 (17.3-24.8) | 1.18 (0.85-1.65) | 1.08 (0.75-1.57) |
| Richer | 16.7 (13.8-20.2) | 1.36 (0.96-1.93) | 1.35 (0.94-1.96) | 21.3 (17.7-25.2) | 0.96 (0.68-1.35) | 0.93 (0.64-1.34) |
| Richest | 14.2 (11.4-17.5) | 1.00 (Ref) | 1.00 (Ref) | 19.0 (15.6-22.9) | 1.00 (Ref) | 1.00 (Ref) |

<sup>a</sup> Sampling weights were used to calculate prevalence
<sup>b</sup> Adjusted for mother's age at birth, ANC visit, maternal undernutrition and parity

### Association with maternal and socioeconomic factors

In unadjusted models, having no ANC visits, and maternal undernutrition were found to be significantly associated with low birth weight for both male and female children. The association for younger maternal age at birth was significant only for male children (Table 2). When we adjusted for socioeconomic variables to see whether they can explain the associations of maternal factors with low birth weight, we found that maternal age at birth <20 years (OR 1.40, 95% CI 1.09-1.78 *vs.* 20-29 years) and maternal undernutrition (OR 1.33, 95% CI 1.05-1.69) were associated with low birth weight in male children. For female children, only maternal undernutrition was found to be significant in adjusted model. The association of no ANC visit with low birth weight was explained by socioeconomic factors (Table 2).

For both male and female children, mother with no education (adjusted OR 2.46, 95% CI 1.49-4.05) and primary education (adjusted OR 1.66, 95% CI 1.06-2.60) had higher likelihood of having low birth weight babies when compared to mothers with higher education, even after adjustment for maternal factors. In compared to households with richest wealth index, children born in households with poorest wealth index were more likely to have low birth weight (adjusted OR 1.58, 95% CI 1.07-2.32 in males, and adjusted OR 1.42, 95% CI 0.98-2.05 in females) (Table 3). There was no association between area of residence and prevalence of low birth weight.

## DISCUSSION

In this study, we found that nearly one in every five children born in Bangladesh had low birth weight and factors like younger maternal age at birth, inadequate ANC visit, and maternal undernutrition were associated with increased prevalence of low birth weight. Also, there were inequalities in low birth weight burden by mother’s education and household’s wealth index.

The prevalence of low birth weight in our study using mother’s perception about baby’s size at birth was about 20%. According to the National Low Birth Weight Survey Bangladesh 2015, the prevalence of low birth weight using the WHO cut-off of <2500 grams was 22% [4,10]. Similar prevalence estimates using these two different methods for defining low birth weight have increased reliability of our findings. That report found that baby girls suffered more from low birth weight than boys, and we also found higher prevalence of low birth weight among girls (22% vs. 18% among boys). Similar findings on sex-difference in low birth weight prevalence was also observed in studies from other low and middle-income countries [11]. Moreover, current literature suggests that the likelihood of mortality during infancy is higher among male babies with low birth weight than female babies with low birth weight [12,13]. Little is known about which factors contributes to the sex-difference in low birth weight. Our results also suggested that the burden of low birth weight in Bangladesh reduced significantly over the last decade – from 30% in 1998 to 20% in 2014 [14]. This reduction in low birth weight has also contributed to reduction of infant mortality in Bangladesh [15].

We found that younger maternal age at birth was significantly associated with low birth weight among boys, even with adjustment for socioeconomic factors. This suggests that the association between them cannot be explained by the confounding effects of socioeconomic factors. We did not observe a significant association for girls, but it’s not clear why such association can differ among boys and girls. Younger mothers tend to be undernourished and develop anemia during pregnancy – which could be the possible mechanisms for this association [16–18]. We also observed significant adverse effects of maternal undernutrition on low birth weight, independent of socioeconomic status. Previous literature also listed smoking, poor diet, deficiency of micronutrients and illness during pregnancy as risk factors of maternal under-nutrition [19,20]. However, the prevalence of smoking among women in Bangladesh is very low and thereby this shouldn’t contribute to low birth weight in Bangladesh [21]. Maternal under-nutrition is related to intra-uterine growth retardation leading to low birth weight. This is because the mother and the fetus compete for nutrition during the pregnancy and under-nutrition in mothers can worsen the situation [19]. In our data, 28% mother was younger than 20 years at the time of their baby’s birth, which explains further needs for preventing child marriage and promoting family planning to the younger women. According to a recent nationwide survey in Bangladesh, adolescent pregnancy was more likely to be associated with infant mortality, low birth weight, and stillbirth in newborns [22].

ANC visit has been playing a pivotal role in reducing maternal mortality, perinatal complications, and low birth weight [23,24]. Our study showed higher odds of low birth weight amongst those who did not have any ANC visit when compared to those who had 4 or more ANC visits. Such association is highly likely to be confounded by lower socioeconomic status because mothers are less likely to have ANC visit and have higher chance of babies with low birth weight. When we adjusted for area residence, household’s wealth index, and mother’s education, the observed association between no ANC visit and low birth weight became attenuated. Studies conducted in Ethiopia and Nepal also found that mothers who had access to ANC during pregnancy had significantly lower risk of delivering a low birth weight baby [25,26]. Some studies also suggested that both adequacy and quality of ANC visits are important to get the full beneficial effects of ANC visit on promoting healthy birth weight [27].

Among mothers with no or primary education, the odds of having low birth weight babies was significantly higher when compared with mothers with higher education. Similarly, a recent study conducted in Malawi reported that mothers with no formal education were more likely to give birth babies with low weight [28]. It is understandable that lower level of education may result in limited access to ANC care as well as low knowledge about healthy diet. It is also difficult for an uneducated mother to adhere to health promotion messages [29–31]. However, in our study we controlled for mother’s age, maternal undernutrition and number of ANC visits, and the association between mother’s education and low birth weight remained significant.

Our study also showed that mothers from poor socioeconomic background had higher odds of having babies with low weight. After adjustment for maternal factors, the association remained significant among boys while it was near to reaching statistical significance among girls. The socioeconomic inequalities in low birth weight was also observed in other settings [11]. Understanding the underlying pathways linking poor wealth and low birth weight is complex, partly because poor social status can affect many factors of a pregnancy which can lead to low birth weight. Some studies suggested that poor mothers could not afford paying for prenatal care services [32,33], but in Bangladesh ANC visit are made free of cost by the government. We did not find any significant difference for low birth weight between rural and urban areas.

One of the limitations of our study is using mother’s perception of child’s size at birth to define low birth weight. However, a recent paper suggested that mother’s perception of birth size is a good proxy for birth weight in large nationally representative surveys [8]. Also, our observed prevalence of low birth weight was similar to that of a recent study in Bangladesh which used measured birth weight to define low birth weight (<2500 grams) [22]. Also, there could be residual confounding due to unmeasured variables as well as inadequate reporting. Therefore, it is important to identify the relative contributions of maternal and socioeconomic factors on low birth weight prevalence with caution - so that effective public health interventions can be taken to curb the burden of low birth weight in Bangladesh. Mother’s education can be an effect measure modifier because self-reported child’s size at birth can vary widely by mother’s education level. As a sensitivity analysis, we looked at the associations among subgroups of women by their education level and found no differences in relevant estimates (data not shown). Prematurity is responsible for large number of low birth weight babies, but we did not have information about gestational age and thereby could not take prematurity into account in our study. The strengths of our study are large sample size, nationally-representative sample, and availability of large number of maternal and socioeconomic variables. Also, to the best of our knowledge our study is the first study in Bangladesh that has looked at the factors for low birth weight separately among boys and girls.

In conclusion, our study highlighted that the prevalence of low birth weight is still high in Bangladesh and higher among girls than boys. We identified that mother’s age less than 20 years, undernutrition, and no ANC visit during pregnancy are associated with higher likelihood of having low birth weight babies. Socioeconomic inequalities in low birth weight burden by mother’s education and household’s wealth index are also observed, but they do not fully explain the associations between maternal factors and low birth weight. Public health interventions should target these areas to promote healthy birth weight in Bangladesh, and beyond.

## AUTHOR STATEMENT

Conception and design: FH, GA and MS

Data collection and management: FH, MS

Data analysis: FH, YA, and MS

Interpretation of the results: all authors

Drafting of the article: FH

Critical revision of the article for important intellectual content: all authors

Final approval of the article: all authors.

## DATA STATEMENT

This study used data from the Bangladesh Demographic and Health Survey 2014, which is available from the DHS program website

